# Distinct dynamic stability patterns among three Atlantic cod subpopulations in the North Sea

**DOI:** 10.1101/2025.07.07.663450

**Authors:** Hsiao-Hang Tao, Manuel Hidalgo, Vasilis Dakos

## Abstract

Atlantic cod (*Gadus morhua*) in the North Sea and adjacent waters has been extensively studied due to its continued population decline despite reduced fishing pressure. Recent evidence reveals that it forms a metapopulation consisting of three subpopulations— Northwestern, Southern, and Viking—highlighting the need to understand their distinct dynamics to support spatially targeted fisheries management. We applied a dynamical systems approach to quantify two key properties for each subpopulation from 1983 to 2023: dynamic stability, indicating whether subpopulations tend to return to or move away from their state each year, and the relative sensitivity of age groups, reflecting the magnitude of change in each group’s abundance in response to perturbations in the age-structured interaction network. We found distinct dynamic stability patterns among the three subpopulations. The Northwestern subpopulation fluctuated between stable and unstable states before stabilizing at low abundance. The Southern subpopulation shifted from high-abundance, stable states toward low abundance at a transient with an uncertain future. The Viking subpopulation remained stable with relatively constant abundance over time. Throughout the study period, the age-1 group remained highly sensitive to perturbations in both the Southern and Viking subpopulations, whereas no single age group showed dominant sensitivity in the Northwestern subpopulation. Fishing mortality causally influenced the abundance and/or stability of the Northwestern and Southern subpopulations, but not the Viking subpopulation. In contrast, sea bottom temperature did not affect the abundance or stability of any of the subpopulations. Our findings reveal distinct patterns in abundance, dynamic stability, relative sensitivity of age groups, and responses to external drivers across the three subpopulations, highlighting the importance of accounting for the complex population structure of Atlantic cod in future research, assessments, and management efforts.

## Introduction

Marine ecosystems provide key functions and services including nutrient cycling, biodiversity, and food resources. Disturbances, such as climate change and overfishing, are seriously threatening the future provision of these services, raising an urgent need for undertaking management actions that promote marine ecosystem stability. The stability of a marine ecosystem is defined as the capacity to persist or maintain its present state in the face of disturbances. When a marine ecosystem is vulnerable, disturbances may destabilize marine populations and the services they provide, making them more sensitive natural and anthropogenic pleasures (Perry *et al*. 2010). Although exploitation rate has declined recently in some fisheries, historically prolonged overexploitation has delayed rebuilding and increased the uncertainty in recovery times in many cases (Neubauer *et al*. 2013). Ocean warming has also caused structural changes in some marine fish communities (Griffith, Strutton and Semmens 2018; McLean *et al*. 2019). Tracking and forecasting changes in the stability of key marine fish populations over time and identify the drivers provides crucial information for managing fisheries toward sustainable harvesting. However, there is a fundamental challenge that prevents the advances in studying marine fish populations.

While most established stability measures assume a stable equilibrium and constant magnitude of disturbances (Donohue *et al*. 2016), marine ecosystems are influenced by multiple disturbances simultaneously, with their magnitude changing with time. Disturbances interact with intrinsic properties (e.g., growth rate of a population) and feedback on each other, so that marine ecosystems usually exhibit state-dependent states (Hsieh *et al*. 2005). State-dependency refers to how the relationship among interacting variables change depending on the system’s current state, leading nonlinear rather than linear dynamics. For example, such nonlinear dynamics is commonly observed in marine fish populations, particularly those that are heavily exploited (Glaser *et al*. 2014). Yet, marine populations are often evaluated based on linear approaches, such as the time to recover to a fixed reference point (Lotze *et al*. 2011). Only recently nonlinear dynamics, discontinuous dynamics, and regime shift dynamics are being emphasized in the study of fisheries (Möllmann *et al*. 2021; Blöcker, Sguotti and Möllmann 2022). Thus, marine ecosystem stability needs to move beyond a static, linear description based on a stable equilibrium towards tracking changes in the stability of marine ecosystems under ever-changing disturbances away from equilibrium.

The Atlantic cod populations in the North Sea and adjacent waters exhibit ecological, economical, and cultural importance. However, the North Sea population has experienced abrupt decline in abundance around year 2000 without a significant sign of recovery (ICES 2024). Recent work examining the nonlinear dynamics of this population has suggested one or multiple shifts in the internal dynamics around year 2000, which were closely related with fishing and warming (Blöcker et al. 2022, 2023). Additionally, a study using dynamic stability approach showed that this population has shifted from a disequilibrium state to a stable, low-abundance state around year 2003 (Tao et al 2025, *in press*).

Yet, recent evidences reveal that Atlantic cod in the North Sea and the adjacent waters has a meta-population structure composed of three subpopulations (ICES 2020) **(Fig 1)**: the Northwestern subpopulation which occupies the northwest of the North Sea and west of Scotland, the Southern subpopulation which distributes in the south-central North Sea, and the Viking subpopulation in the northeast of North Sea. These three subpopulations differ in genetic structure, morphology, demographic traits, physiology (e.g., thermal tolerance), and phenotypic traits (e.g., rates of maturity and growth (ICES 2020) so that since 2024 stock assessments are conducted separately for each subpopulation (ICES 2024). This new information motivates further research into the dynamics and resilience at the subpopulation level to support more targeted fisheries management. For example, recent studies have shown that three subpopulations of Atlantic cod exhibit distinct regime shift dynamics and that the effects of fishing and sea temperature differed across them (Cecapolli *et al*. 2025). What remains unclear is how the dynamic stability of these three subpopulations has changed over time, and what are the drivers of the dynamic stability of each subpopulation.

**Fig. 1.**
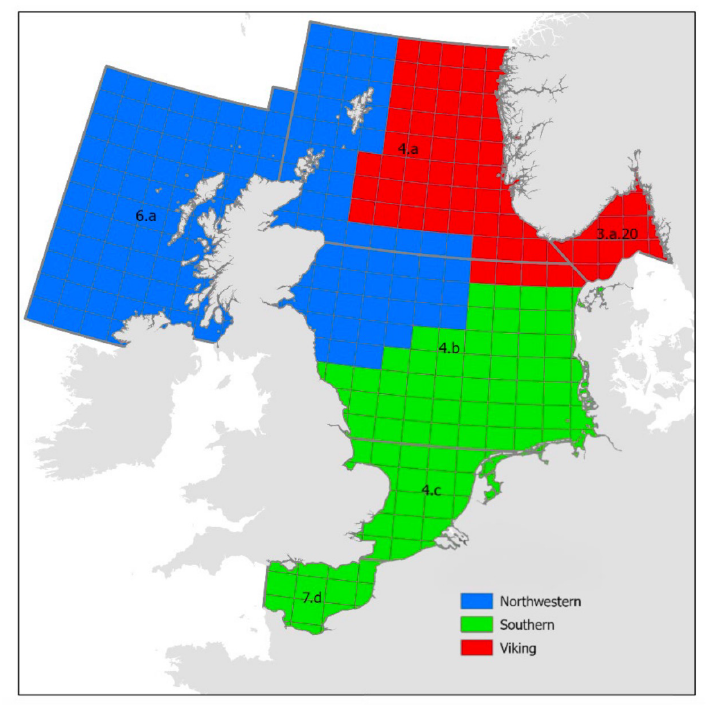
Distribution of three subpopulations of Atlantic cod in the North Sea. (source : ICES 2024)

In this work, we used empirical dynamic modelling to study the dynamic stability and drivers of three subpopulations of Atlantic cod in the North Sea and adjacent waters between 1983-2023. Using assessed abundance and fishing mortality (ICES 2024) as well as sea bottom temperature measured during over four decades of surveys (ICES data portal), we address three research questions: 1) what is the pattern of dynamic stability of each subpopulation, 2) what is the relative sensitivity of age groups in each subpopulation; 3) and what are the drivers of the dynamic stability and abundance of each subpopulation.

## Materials and Methods

We obtained the estimated yearly abundance of each subpopulation and fishing mortality from the latest stock assessment model of the three subpopulations of Atlantic cod in the North sea and adjacent waters between 1983-2023 (ICES 2024). This model assessed three subpopulations separately, using fisheries-dependent information and the IBTS winter survey data including the substocks of cod in the North Sea (Subarea 4), West of Scotland (Division 6.a), the Skagerrak (Subdivision 20), and the eastern English Channel (Division 7.d). In the assessment model, it is assumed that there are mixings between juveniles of Viking and Southern subpopulations (ICES 2024). Mean fishing mortality between 2-4 years was used to indicate fishing pressure as they are the most targeted age groups.

The yearly winter sea bottom temperature between 1983 to 2023 was calculated based on the bottle and low-resolution CTD data (assessed from the ICES data portal https://data.ices.dk/view-map). As Atlantic cod mostly live up to 200 m depth, we extracted the winter temperature recordings (from January to March) from all cruises and stations between the sea surface and 200 meters within the distribution area of each subpopulation and calculated the mean yearly temperature. We used the winter data because the estimated cod abundance from the stock assessment is based on the winter survey data.

To estimate the stability and relative sensitivity for each subpopulation, we estimated the optimal embedding dimension of each age group by Simplex Projection (Sugihara and May 1990) using the normalized abundance time series of each age group. Subsequently, we examined the nonlinearity of each subpopulation using S-map (Sugihara 1994). Then, we used MDR S-map (Chang *et al*. 2021) to reconstruct the state space and obtain the pair-wise age group interaction matrices for each year. From each interaction matrix, we extracted the dominant eigenvalue and dominant eigenvectors to calculate the dynamic stability and relative sensitivity between age groups, respectively. For a detailed analytical framework see (Tao et al 2025, *in press*).

The dynamic stability is the absolute value of the dominant eigenvalue from the age interaction matrix, indicated as |DEV| (Ushio *et al*. 2018). The dynamic stability refers to whether the system is converging to (when |DEV| < 1; we refer to locally stable) or diverging from (when |DEV| > 1; we refer to locally unstable) the trajectory, when given a small perturbation (Ushio *et al*. 2018). When |DEV| = 1, the system is neither converging nor diverging, meaning that it is at the threshold between stability and instability.

The biological meaning of dynamic stability can be often confusing. A stable system does not mean that the system is necessarily more desirable, more productive or in a better sustainable state than an unstable system. Instead, a dynamically stable system based on the definition we use here (|DEV| < 1) simply means that the system is likely to stay close to its current state. For example, a population at the high-abundant stable state implies it is productive, while a population at low-abundant stable state implies that it is trapped in a low-productivity state without a change in the near future. Similarly, a system which is dynamically unstable (|DEV| > 1) does not necessary imply that it is at risk, but rather indicates a potential change is going to happen to the system state, although the direction of change is unknown.

We also estimated the relative sensitivity of age groups as the absolute value of the dominant eigenvector of each age group from the age interaction matrix derived from the MDR S-map, after being standardized by the square root of sum of squares of all eigenvectors. The relative sensitivity refers to the distance between unperturbed abundance at a time point and perturbed abundance at the next time point (Medeiros *et al*. 2022). That is, a relatively sensitive age group has relative larger changes in its abundance after a perturbation to the abundance of all age groups.

Lastly, to examine the drivers of the dynamics for each subpopulation, we performed convergent cross mapping (ccm) (Sugihara *et al*. 2012) to test for causal relationships between potential drivers (fishing mortality and sea bottom temperature) and subpopulation dynamics (dynamic stability and abundance). For each pair of variables, we first applied ccm to the original time series, using 5,000 bootstrapped subsets of time points. We then repeated the analysis using 5,000 bootstraps of randomly shuffled time series of the variables as a null model to estimate the expected level of forecasting skill in the absence of causality. Causality was considered significant if both of the following criteria were met: i) the forecasting skill (rho) increased monotonically with increasing time series of the response variable (i.e., Mann-Kendall’s test: τ > 0, *p* < 0.05); and ii) the median rho from the original bootstrapped time series exceeded the 95th percentile of the rho distribution from the null model.

## Results

### 1. Abundance and dynamic stability of subpopulations

By combining abundance trends with dynamic stability, we observed that the three subpopulations exhibited distinct internal dynamics between 1983 and 2023. The Northwestern subpopulation transitioned from a high-abundance, fluctuating state to a low-abundance, stable state. In the first half of the study period, abundance fluctuated substantially, while dynamic stability alternated between stable and unstable states (**Fig. 2a– b**). In the latter half, abundance declined to around 250,000, and dynamic stability remained predominantly stable. These patterns suggest that the Northwestern subpopulation has stabilized at a low-abundance state and is unlikely to undergo significant changes in the near future.

**Fig. 2.**
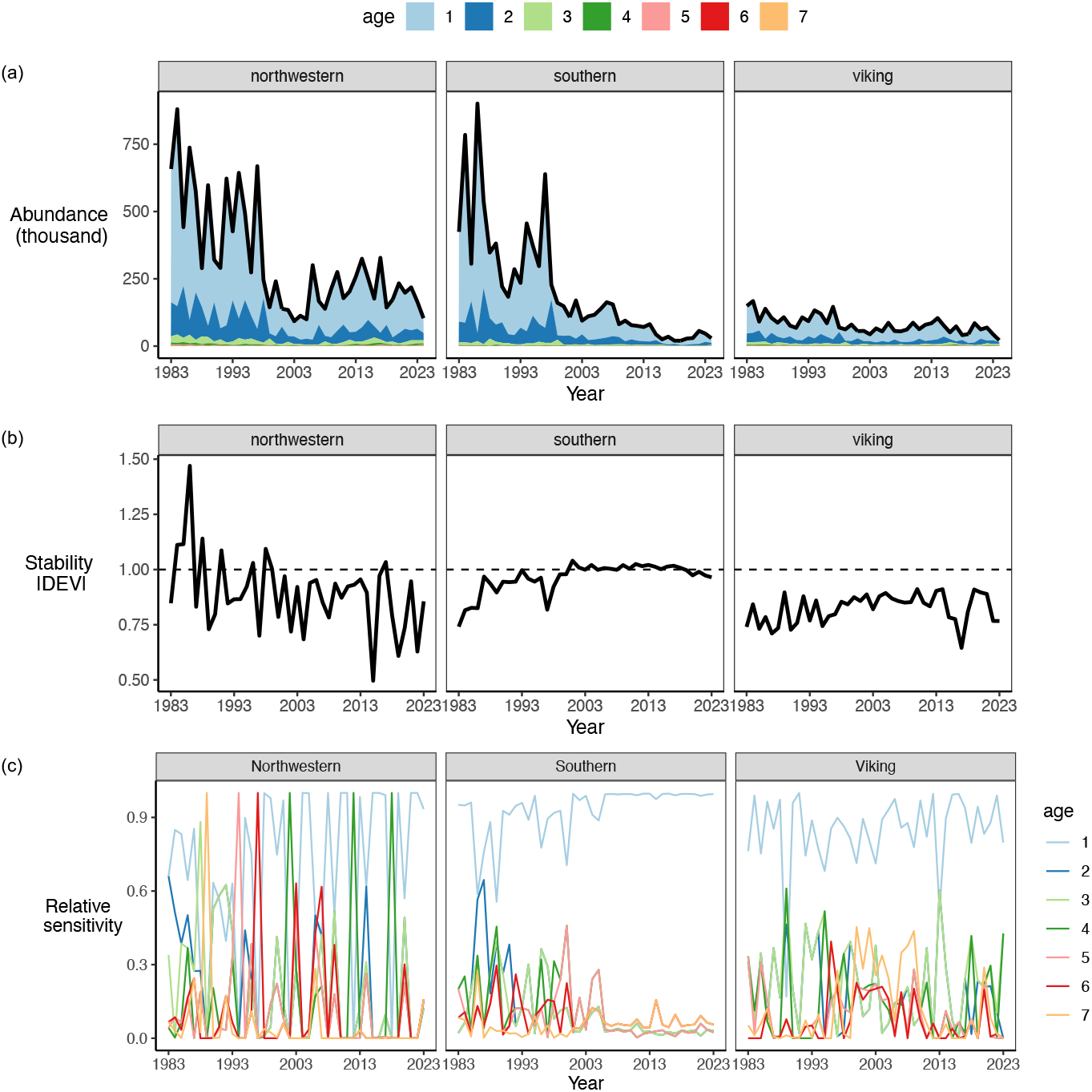
Abundance, age structure, dynamic stability, and relative sensitivity of age groups for each subpopulation. (a) Estimated abundance as stacked plots with color indicating age groups. (b) Stability indicated by |DEV| derived reconstructed age-group interaction matrices using MDR S-map. Dotted line indicates the threshold between stability and instability at |DEV| = 1. |DEV| > 1 indicates the system is locally unstable while |DEV| < 1 indicates the system is locally stable. (c) Relative sensitivity of age groups derived as standardized absolute value of eigenvector of reconstructed age-group interaction matrices. Colors indicate age groups.

In contrast to the Northwestern subpopulation, the Southern subpopulation transitioned from a fluctuating, high-abundance stable state to a low-abundance state. Specifically, in the first half of the study period, this subpopulation exhibited large fluctuations in abundance and gradually shifting from stable states approaching the threshold between stability and instability (**Fig 2a-b**). In the second half of the study period, the abundance dropped abruptly while |DEV| remained very close to the threshold between stability and instability, but on the unstable side. This suggests that this subpopulation stayed at a transient (Hastings *et al*. 2018), moving towards a new stable or unstable state in the near future.

The Viking subpopulation remained stable at a relatively constant abundance throughout the whole study period (**Fig 2a-b**), indicating a strong tendency to persist in this state in the near future.

### 2. Relative sensitivity of age groups

The relative sensitivity of age groups varied among the three subpopulations and may explain differences in their dynamic stability patterns (**Fig 2b-c**). In the Northwestern subpopulation, the relative sensitivity of age groups changed over time without a consistently dominant sensitive age group. This variability could be linked to the highly fluctuating dynamic stability of this subpopulation. In contrast, in the Southern subpopulation, age-1 group remained the most sensitive among all throughout the period, with sensitivity peaked during the latter half of the study period. This stability in sensitivity corresponds to the dynamic stability remaining close to the threshold between stability and instability at this time. Age-1 group was the most sensitive in the Viking subpopulation throughout the study period, and its fluctuations in sensitivity closely mirrored the overall dynamic stability of the Viking subpopulation.

### 3. Drivers of abundance and stability of subpopulations

Fishing mortality for all three subpopulations remained above the FMSY reference level throughout the study period, although it began to decline around the year 2000 (**Fig. 3a**). The Southern subpopulation experienced the highest fishing mortality, followed by the Northwestern and then the Viking subpopulations.

**Fig 3.**
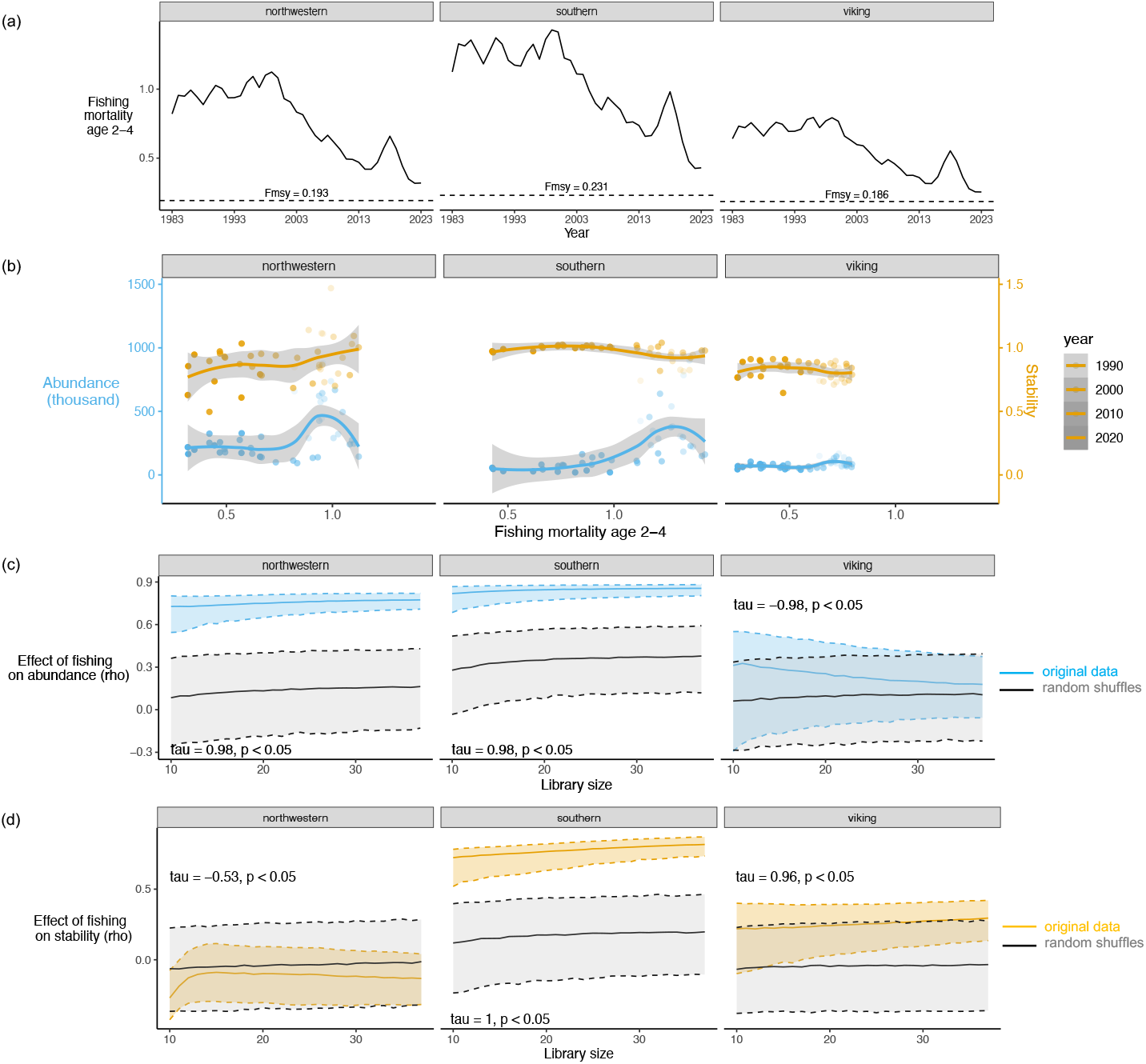
Relationship between fishing mortality with abundance and stability of the three North Sea cod subpopulations. (a) Temporal changes in the mean fishing mortality of 2-4 years between 1983-2023. Dotted horizontal line shows the fishing mortality at maximum sustainable yield (F_MSY_) reported in (ICES 2024). (b) Correlations between fishing mortality with abundance (blue) and stability (yellow). The fitted lines are loess smooth curves. (c) Causal effect of fishing on abundance. The forecasting skill (rho) from the convergent cross mapping was examined across library sizes. The blue lines and shaded areas represent the median and 5–95% quantiles of the forecast skill from 5000 bootstraps of real time series. The grey line and shaded areas represent the mean and 5–95% quantiles of 5000 shuffled time series. The causality is significant if i) rho showed a monotonic increase with library size (i.e., when Mann-Kendall’s test shows tau >0 and *p* < 0.05); and if ii) the median forecasting skill from real time series bootstraps exceeded the 95% quantile of shuffled time series.

Fishing mortality causally influenced the abundance of the Northwestern and Southern subpopulations **(Fig. 3c, S1)**. In addition, fishing mortality causally influenced the stability of the Southern subpopulation but not that of the Northwestern subpopulation **(Fig. 3d, S1)**. No causal links were found between fishing mortality and either abundance or stability in the Viking subpopulation **(Fig. 3c–d, S1)**, likely reflecting the lower fishing pressure it experienced **(Fig. 3a)**. Importantly, despite some causal relationships, there was no apparent correlation pattern between fishing mortality and abundance or dynamic stability in any subpopulation **(Fig. 3b, S1)**.

No causal relationships were found between sea bottom temperature and either abundance or dynamic stability in any subpopulation (**Fig. 4c**), despite negative correlations with abundance observed for the Northwestern and Southern subpopulations (**Fig. 4b, S1**).

**Fig 4.**
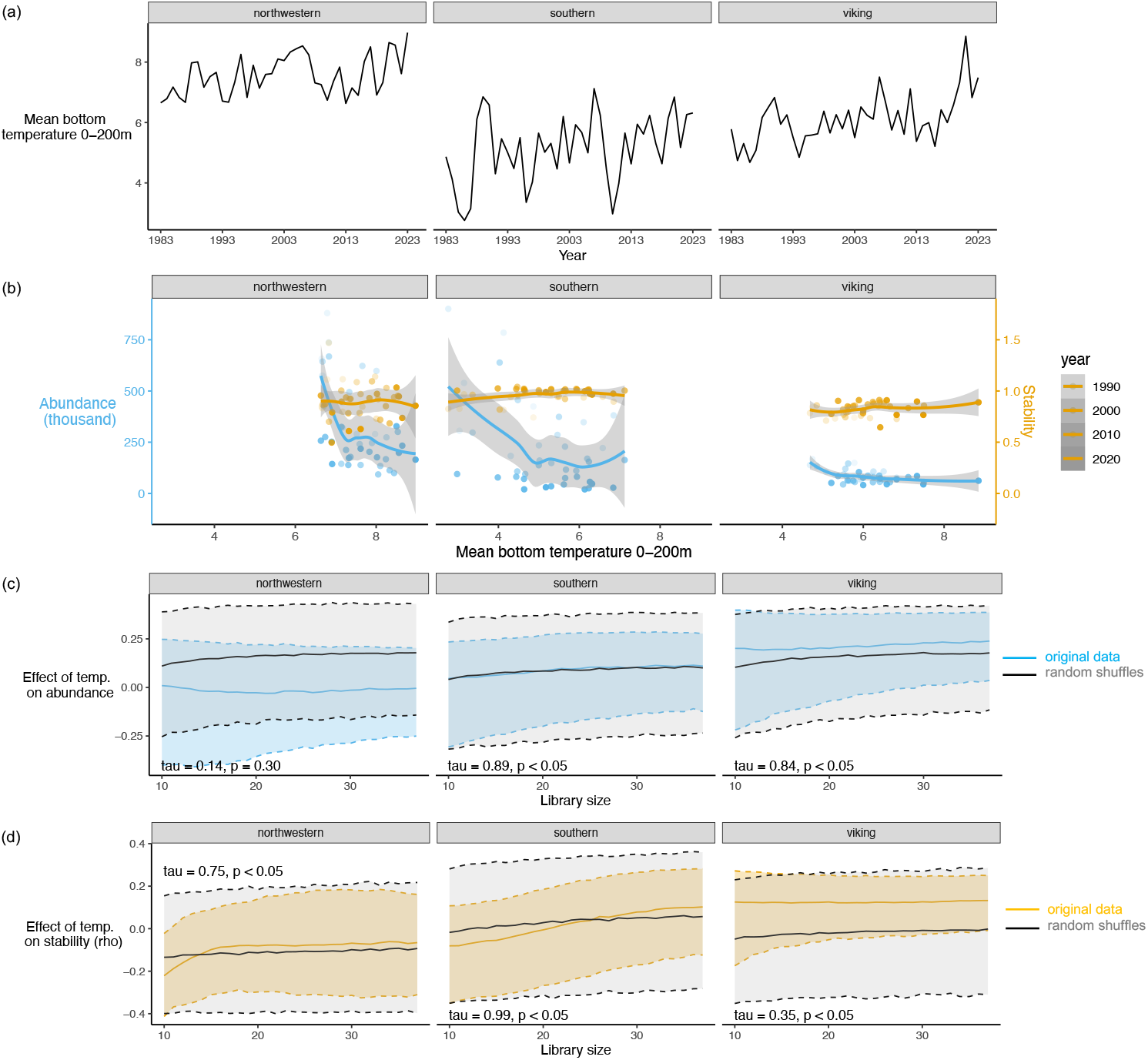
Relationship between sea bottom temperature with abundance and stability of the three North Sea cod subpopulations. (a) Temporal changes in the mean sea temperature of 0-200m between 1983-2023. Dotted horizontal line shows the fishing mortality at maximum sustainable yield (F_MSY_) reported in (ICES 2024) (b) Correlations between fishing mortality with abundance (blue) and stability (yellow). The fitted lines are loess smooth curves. (c) Causal effect of fishing on abundance. The forecasting skill (rho) from the convergent cross mapping was examined across library sizes. The blue lines and shaded areas represent the median and 5–95% quantiles of the forecast skill from 5000 bootstraps of real time series. The grey line and shaded areas represent the mean and 5–95% quantiles of 5000 shuffled time series. The causality is significant if i) rho showed a monotonic increase with library size (i.e., when Mann-Kendall’s test shows tau >0 and *p* < 0.05); and if ii) the median forecasting skill from real time series bootstraps exceeded the 95% quantile of shuffled time series.

## Discussion

While tracking changes in the stability of marine fish populations over time is useful for fisheries management toward sustainable harvesting, many marine fish populations exhibit nonlinear dynamics (Glaser *et al*. 2014) and requires methods beyond linear approaches. We used a dynamical systems approach to estimate the dynamic stability of three subpopulations of Atlantic cod in the North Sea. We found that the three subpopulations exhibit different dynamical stability patterns. Strikingly, even the Northwestern and Southern subpopulations display similar abundance trends, they differ markedly in their dynamic stability patterns. These evidences suggest that measuring dynamic stability of natural populations offers insights for characterizing and assessing beyond their productivity status, and thus complementary to biomass or abundance dynamics.

Our results based on dynamic stability reveal both consistencies and discrepancies when compared to a recent study on regime shifts in three subpopulations of North Sea cod (Cecapolli *et al*. 2025). One key similarity is that while our result showed that while the Southern subpopulation has remained at a transient state since the 2000s (**Fig. 2b**), Cecapolli et al. (2025) reported that this subpopulation underwent a regime shift and kept moving in and out from the bifurcation zone. Another similarity is that both works suggest increasing stability of the Northwestern subpopulation in recent years. Our findings show that while this subpopulation fluctuated between stable and unstable states in earlier years, it has remained predominantly stable after year 2000. Cecapolli et al. (2025) also identified that this subpopulation approached a tipping point in the 1990s but was subsequently pushed into a stable state, driven by changes in fishing efforts and sea surface temperature.

However, a notable discrepancy arises in the interpretation of the Viking subpopulation. Our results indicate that this subpopulation remained stable throughout the study period, whereas Cecapolli et al. (2025) concluded that it crossed a tipping point around 2000 before eventually reaching a stable state. This difference may be attributed to the distinct methodologies used in these two studies. On one hand, the dynamic stability estimated in our study does not assume equilibrium and does not require prior knowledge of the stability landscape. Moreover, dynamic stability is derived solely from interactions among age groups, without explicitly accounting for external drivers such as environmental conditions or fishing pressure (Ushio *et al*. 2018). On the other hand, the regime shift analysis used in (Cecapolli *et al*. 2025) identifies transitions between alternative system states by examining changes in the relationships between system states (e.g., abundance, biomass) and external drivers (e.g., Blöcker et al. 2023). While these methodological differences likely explain the divergent interpretations of the Viking subpopulation’s dynamic stability, both studies reached consistent conclusions regarding the Northwestern and Southern subpopulations. Altogether, these results suggest that measuring and comparing various available resilience indicators can offer a comprehensive understanding of population status.

The seemingly similar patterns between the sensitivity of the age-1 group and dynamic stability in both the Southern and Viking subpopulations may be explained by two main factors: (1) the relatively high abundance of the age-1 group compared to other age classes,and (2) the strong dependence of age-1 juveniles on the reproductive output of mature adults. The second factor is particularly relevant to the Southern subpopulation, where a decline in adult abundance during the latter half of the study period may have reduced juvenile recruitment, thereby increasing the sensitivity of the age-1 group to fluctuations in other age groups. In contrast, this explanation does not apply to the Viking subpopulation, where the age structure remained relatively stable. Instead, the high sensitivity of the age-1 group in this subpopulation is more likely driven by its consistently higher abundance compared to other age groups.

Fishing pressure has a stronger impact on the Southern subpopulation compared to others, as it causally influences both its abundance and dynamical stability. This finding is consistent with previous evidence showing that fishing activity has historically been more intense for Atlantic cod in the Southern North Sea (Engelhard, Righton and Pinnegar 2014). However, no apparent correlation was found between fishing mortality and either abundance or dynamical stability across any subpopulation **(Fig. 3b, S1)**. This inconsistency is likely due to variations in these correlations across different regime shift periods, reflecting underlying nonlinear relationships between the variables, as illustrated by Cecapolli *et al*. (2025).

In contrast to fishing pressure, we found no causal links between winter sea bottom temperature and either abundance or dynamic stability in any subpopulation—even though negative correlations with abundance were observed in the Northwestern and Southern subpopulations (**Fig 4**). This finding implies that, in natural systems, correlation does not necessarily imply causation (Sugihara *et al*. 2012). However, it is important to note that we used winter sea bottom temperature rather than annual sea bottom temperature, as the abundance data used in this study were estimated based on winter survey observations (ICES 2024: 202). In contrast, earlier studies at both the population and subpopulation levels of North Sea cod have demonstrated the influence of environmental drivers. For instance, Cecapolli *et al*. (2025) reported that both annual sea surface temperature and sea bottom temperature affect the surveyed abundance of three subpopulations, with the strength of these influences varying among them. Additionally, Engelhard, Righton and Pinnegar (2014) showed that annual sea surface temperature has driven a northward spatial shift of the North Sea cod population.

This study has several methodological limitations. First, dynamic stability is often underestimated due to process and measurement errors, yet our interpretation relies heavily on whether dynamic stability is above or below the threshold when |DEV| =1. This introduces uncertainty into our results and conclusions. Second, validating results from empirical dynamic modeling is challenging because of the inherent complexity of nonlinear population dynamics. Third, while an unstable system state indicates a likely change in the near future, the direction of that change—whether abundance or biomass will increase (recovery) or decrease (further decline)—remains unknown. Forth, the subpopulations have been treated as isolated units, while early life stage connectivity and adult movement have not been considered as a potential driver of dynamic stability (ICES 2020). Addressing these limitations through methodological advances will improve the reliability of results and their usefulness for precautionary management.

Finally, we highlight that there is growing relevance of developing and applying nonlinear dynamics-based methods in assessing natural population and communities beyond marine ecosystems (Cenci and Saavedra 2019; Medeiros and Saavedra 2023; Zhao *et al*. 2023). Nonlinearity is common in natural populations including insects, bony fish, mammals, and birds (Clark and Luis 2019), and such nonlinearity can be reinforced by environmental change and human activity (Glaser *et al*. 2014). These evidences underscore the need to using nonlinear methods that track the resilience of natural populations under ever-changing disturbances away from equilibrium.

## Data and code availability

Estimated abundance of three subpopulations and fishing mortality data are obtained from (ICES 2024). Sea bottom temperature data is accessed from ICES data portal https://www.ices.dk/data/data-portals/pages/default.aspx. We used “rEDM” package (https://cran.r-project.org/src/contrib/Archive/rEDM, version 1.2.3) to perform all the analysis. The MDR S-map was performed using functions provided in Chang et al. (2021). All statistical analyses were performed in R 4.3.1. Codes for reproducing the results are deposited in https://github.com/HsiaoHang/git_cod_stability_subpopulation.

## Acknowledgements

We thank the research and survey teams of International Bottom Trawl Survey from International Council for the Exploration of the Sea (ICES) and Working Group on the Assessment of Demersal Stocks in the North Sea and Skagerrak (WGNSSK) for providing the data. We thank Chih-hao Hsieh, Alessandro Orio and Masayuki Ushio for helpful discussions. This work is funded by Marie Curie postdoctoral fellowship of H.H-T. (HORIZON-MSCA-2021-PF-01, European Commission, LSP 244267; MAFIS project). MH acknowledges within CORES project (PID2023-151282OB-I00, Spanish Ministry of Science and Innovation, MCIN/ AEI /10.13039/501100011033).

## Author contribution

All authors designed the original study, H.H-T. performed the analysis, all authors discussed and interpreted the data, H.H-T. wrote the first draft of the manuscript, all authors contributed substantially to the final version of the manuscript.

