## Supplementary material for "Distinct dynamic stability patterns among three Atlantic cod subpopulations in the North Sea": S1

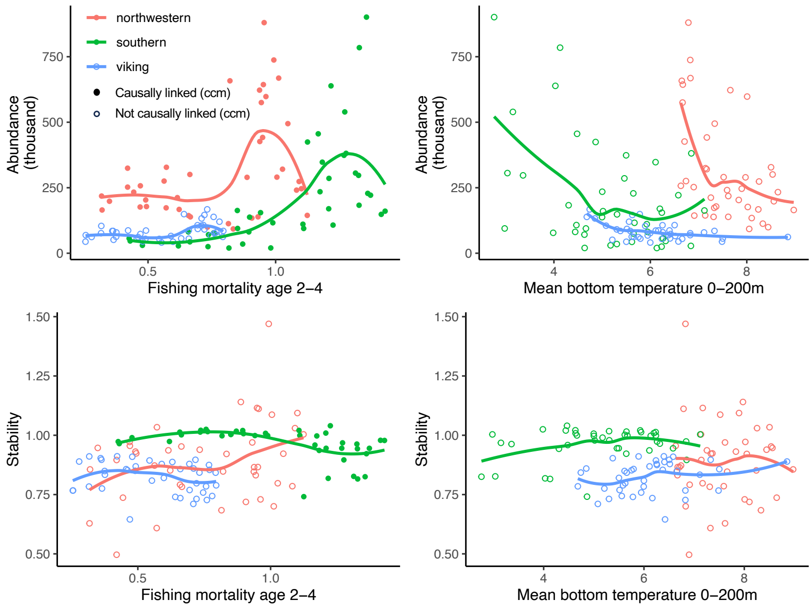


**Fig S1 Relationships between abundance or stability with fishing mortality or mean bottom temperature.** Colors indicate subpopulations. Points indicate yearly values. Fitted lines are loess smooth curves. Filled and open points indicate significant and insignificant causal relationships tested using convergent cross mapping.
